# A candidate electromagnetic channel for coordination between physically separated Drosophila

**DOI:** 10.64898/2026.07.23.740082

**Authors:** Katerina Douka, Efthimios M. C. Skoulakis, Luca Turin

## Abstract

While filming Drosophila melanogaster recovering together from xenon anesthesia, we noticed frequent synchronous pauses in flies recovering in the same chamber. To ask whether such coordination could survive physical separation of the animals, we housed flies individually in the opaque wells of a multi-well plate, with no visual or tactile contact. They still coordinated the magnitude of their movements around a shared light*dark startle more closely than the shared stimulus alone can explain. We isolate this excess coordination with a measure that subtracts the stimulus-driven component by comparing movements within versus across transitions. A complementary common-mode statistic shows that flies also remain weakly but reproducibly correlated away from the transitions. The tested conventional channels do not readily account for the effect: vision is excluded by the opaque walls, and self-generated substrate vibration is disfavored for the background common mode because coordination does not scale with movement amplitude and persists when the continuous side-wall path between wells is interrupted by air gaps. The transition signal declined across the material series from plastic toward aluminum. This pattern is not a simple metal-versus-nonmetal split. It is consistent with material-dependent near-field magnetic radiofrequency coupling through the well geometry and transparent imaging aperture. Vector-network measurements also indicate that near-field inter-well coupling differs between steel and aluminum. We have previously shown that Drosophila emit spontaneous, metabolically powered magnetic radiofrequency radiation that tracks nervous-system activity. Together these results are consistent with a near-field magnetic emission that can leak between separated individuals. We discuss radical-pair detection, possible emission mechanisms, and experiments needed to identify the carrier frequency, detector, and biological role.

**Significance:** Fruit flies housed in separate, opaque chambers that block sight and touch nonetheless coordinate their movements. The coordination persists when mechanical vibration paths between chambers are cut. The coordination is absent when the plate is made of aluminum, and intermediate in size when the plate is made of ferromagnetic steel. This ordering, together with direct RF-coupling measurements, points to a near-field magnetic radiofrequency signal passing between individuals, though the carrier frequency and detector remain unknown. Because flies are already known to emit faint, metabolically powered radiofrequency fields that track nervous-system activity, these results raise the possibility that nervous systems both emit and detect such fields. If confirmed, this would identify an unrecognized magnetic radiofrequency channel in animal biology and suggest that brains are less electromagnetically private than assumed.

## Introduction

The observation that prompted this work came from experiments on the recovery of Drosophila melanogaster from xenon anesthesia ^1^. Forty flies were loaded and anesthetized together inside a polycarbonate syringe used as a hyperbaric exposure vessel; approximately 35 were successfully tracked per video during decompression, with movement quantified frame by frame as the number of pixels changing between consecutive frames. Recovery from a general anesthetic should reflect individual variation in drug sensitivity, body size, and metabolic state, so flies recovering together should resume movement asynchronously. Instead, at 30 fps we observed frequent frames in which every fly was motionless while most were moving in the frames on either side. That shared-chamber observation is retained as the origin of the question (Figure S1), but the evidence analyzed below comes from the isolated-well experiments, where visual and tactile contact were removed.

This pointed to coordination between flies, but flies sharing an enclosure can interact through several channels—vibration, chemical signaling, tactile contact, or vision—any of which could account for it. To test whether the coordination survived physical separation of the flies and to eliminate as many of these confounds as possible, we housed each fly individually while still monitoring them all simultaneously. We used the Zantiks ^2^ multi-well behavioral system, which records the locomotor activity of individual flies in the separate wells of a 48-well plate under programmable light/dark protocols, with automated tracking at 1-second resolution. Each fly was housed in its own well, with 0.5 mL of food consisting of agar and sugar at the bottom. Opaque 3D-printed plates remove visual contact. With the flies thus isolated, we expected the coordination to disappear. It did not. Here we investigate which sensory channel could allow physically separated flies to coordinate their movements.

We report two behavioral phenomena: a fast transition signal locked to the light-to-dark startle, and a weaker background common mode between startles. We then ask whether visual contact, tactile contact, substrate vibration, plate material, and direct RF-coupling measurements fit these observations. The results are consistent with near-field magnetic RF coupling, but they do not by themselves identify the emitter, detector, or carrier frequency.

## Results

Flies left isolated and unstimulated in their wells quickly become quiescent, and coordination cannot be measured in animals that are not moving. We therefore used a one-hour protocol of interleaved light and dark periods of one minute each, giving 30 transitions of each type. Each light*dark transition produces a sharp, reproducible startle response across the whole plate ^2^. It seemed interesting to ask whether flies coordinated their activities after a startle, and whether coordination extended to periods without stimulus. We used two complementary analyses: transition-window excess coordination (Δ) and background common-mode coordination (χ), schematized in Figure 1 and defined in Methods.

**Figure 1.**
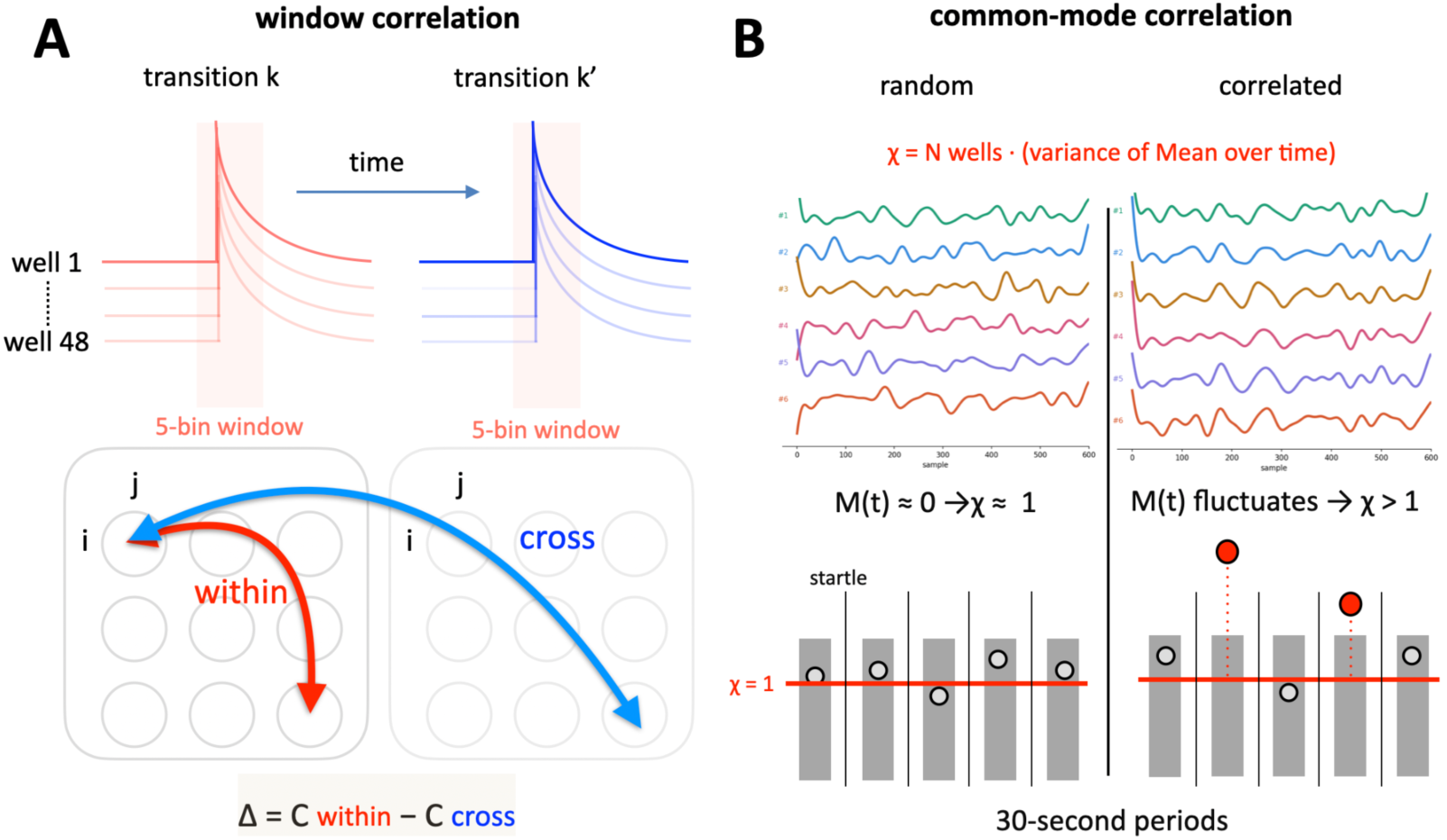
Two complementary measures of inter-well coordination. (A) Transition-locked excess coordination, Δ. Around each light-to-dark switch, the shared light response makes wells correlated even without fly-to-fly coupling. For each pair of wells we therefore compare correlations within the same transition with correlations for the same wells across different transitions. The cross-transition comparison preserves the stimulus response but breaks same-moment coordination. Δ = within – cross is plotted in sliding windows around the switch and is positive when flies sharing a plate move together more than the light cue alone predicts. (B) Background common-mode coordination, χ. Away from transitions, each well is standardized within the central window of a light or dark period and the population mean M(t) is formed across wells. Independent well activity cancels in this average; shared fluctuations remain. χ = N·Var(M) has expectation 1 for independent wells and rises with the mean pairwise correlation φ as χ = 1 + (N – 1)φ. Thus Δ is specific to fast transition-locked coordination after subtracting the shared startle, whereas χ is a sensitive but less source-specific measure of slow shared activity across the plate.

The wells are separated by solid 3D-printed plastic, so there is no visual contact between flies, and the walls rule out tactile contact.

### The light-to-dark switch reveals excess coordination

The light-to-dark switch makes otherwise quiet flies move in synchrony when startled. The question is whether, during a given light-dark transition, flies in the same plate resemble one another more than expected from the shared startle alone. We used the repeated transitions to subtract the shared startle. For each pair of flies, we compared correlations within the same transition with correlations between different transitions. This statistic separates genuine inter-fly coordination from the shared startle response to each light transition. A 5-bin (5 s) window is stepped across each transition, and within it every fly’s activity pattern is correlated with every other’s. Because the correlation spans several bins, it captures the temporal pattern of each response rather than a single instant. Correlations are computed both for flies exposed to the same transition and for flies exposed to different transitions, and only their difference is retained. The stereotyped startle, being reproduced at every transition, contributes equally to both and can be subtracted; what remains is the component two flies share on a given transition but do not reproduce across transitions, which constitutes coordination beyond the common stimulus. A positive Δ means that the plate contains more coordinated movement than the common stimulus explains. Because each point averages a 5-bin (5 s) window, a window centered just before the transition already contains the startle, which produces an apparent ≈2-bin anticipation.

In opaque white plastic plates, Δ rose sharply above startle at the light-to-dark switch in all three movement measures (Figure 3). Dark-to-light transitions gave a smaller signal (Figure 8). The two magnitude measures, DIST (distance travelled) and MSD (mean-squared pixel change), gave the broadest and clearest peaks, reaching about 0.03 around the switch, while the binary moved/not-moved signal was narrower. Thus the effect is not just that flies move at the same time; the amount they move is coordinated too, and movement magnitude carries the cleaner signal. This is the central surprise of this paper: flies that cannot see or touch one another nevertheless move together, raising the question of what physical channel connects separated wells.

**Figure 2.**
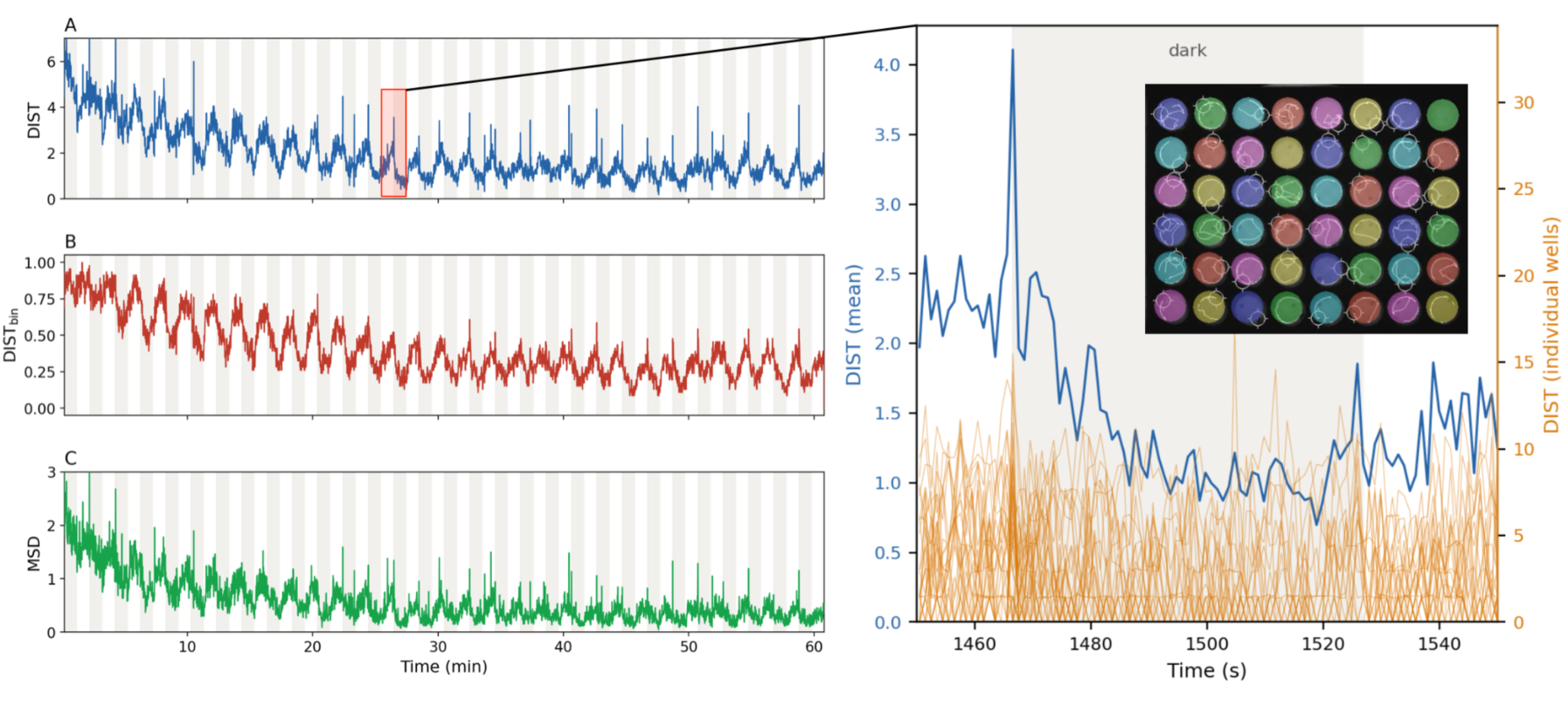
Typical locomotor data and the three movement signals. Activity of flies in a 48-well plate (video image in inset at right) over the one-hour protocol of alternating one-minute light and dark periods (light/ dark blocks shaded), showing the sharp, reproducible startle at each light*dark transition. The traces represent the average of all 48 wells. Left panel: A, B, and C are the three quantities derived from the tracking output and used throughout: displacement (DIST), the distance moved per time bin; its binarized form DIST_bin (moved/did-not-move); and mean-squared pixel change (MSD). Inset: a single DIST transition expanded.

**Figure 3.**
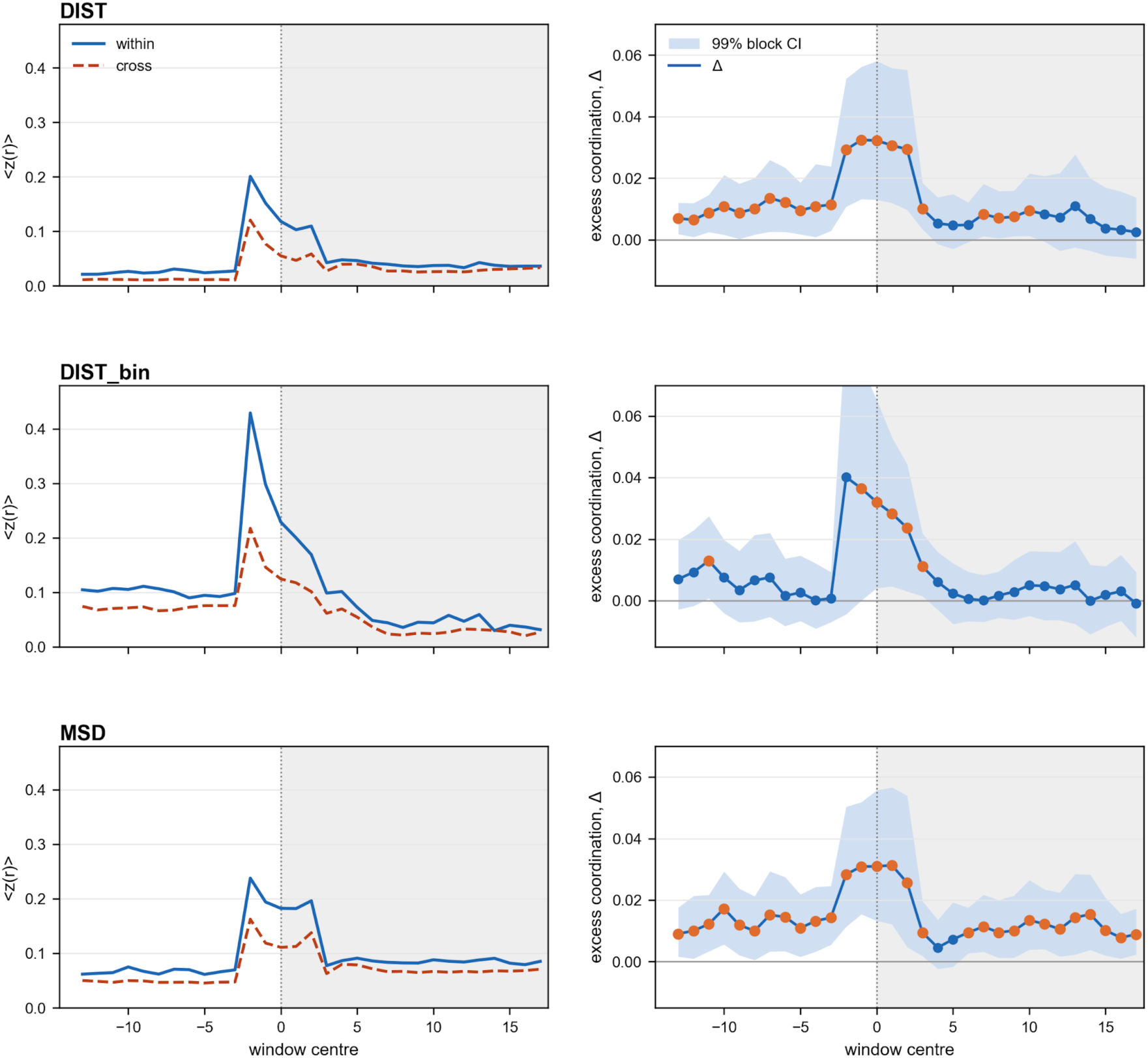
Sliding-window coordination at the light-to-dark transition in white plastic plates. For each signal type (DIST, DIST_bin, and MSD), activity was analysed in 5-bin (5 s) windows whose centres span –15 to +15 bins relative to the light-to-dark transition. Left panels show Fisher-z-transformed within-transition and cross-transition pairwise correlations. Right panels show excess coordination, Δ = within – cross. Shaded bands are 99% moving-block bootstrap confidence intervals from 10,000 transition resamples; filled orange points mark windows whose 99% CI excludes zero. Grey shading marks the dark phase after the transition. Data are from 11 white/plastic plates, comprising 319 light-to-dark transitions. The peak appears slightly before zero because each point averages a 5-bin (5 s) window. A window whose centre lies just before the transition therefore already contains the transition, so the startle contributes before the plotted centre reaches zero.

### Background common-mode correlation

Away from the switches, does fly movement in the whole plate rise and fall together? To measure this, we looked at the middle 30 s of each light and dark period, when no transition was occurring. In each window we standardized each well and averaged the wells at each time point. If flies are independent, this plate average should stay small and χ = N·Var(M) should be near 1. Values above 1 mean that the wells share a common fluctuation. The equivalent pairwise correlation, φ, is small, but χ makes it visible by summing the weak shared signal over many wells. The circular-shift test is described in Methods.

By this measure, flies are not independent even between startles. Figure 4 isolates the data from opaque plastic plates because these plates provide the cleanest test of the background common-mode signal in the absence of metal shielding. In plastic, χ was intermittently elevated above the circular-shift surrogate band in both dark and light periods, and the corresponding significant windows had positive pairwise φ. The statistic is defined in Methods.

**Figure 4.**
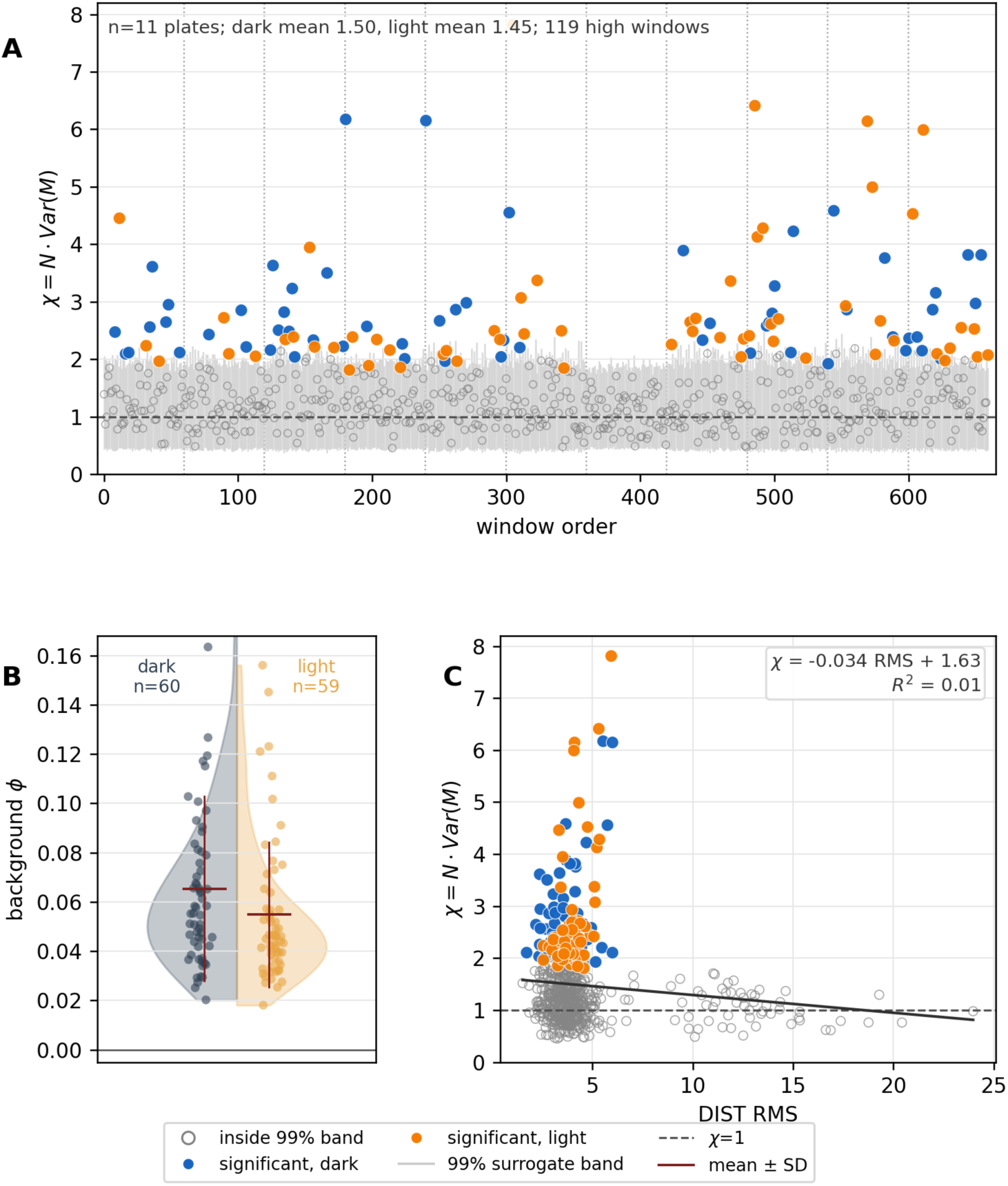
Background common-mode coordination in opaque white plastic plates. (A) DIST χ for the central 30 s of each light or dark period in 11 plastic plates, ordered by plate and time. Open gray circles are windows inside the 99% circular-shift surrogate band; vertical gray bars show the band; blue and orange points mark dark and light windows above it. The dashed line marks χ = 1, the value expected without shared movement. Pooled window means are χ = 1.50 in dark and 1.45 in light; averaging both phases within each plate gives 1.48 ± 0.20 across plates. (B) For the 119 high-χ windows in A, background pairwise correlation φ is shown separately for dark and light periods. Points are individual windows, half violins show the distributions, and dark-red bars show mean ± SD. (C) χ for all windows from A plotted against DIST root mean square (RMS). The fitted line is nearly flat (R² = 0.01), so elevated χ is not explained by large movement bouts alone.

**Figure 5.**
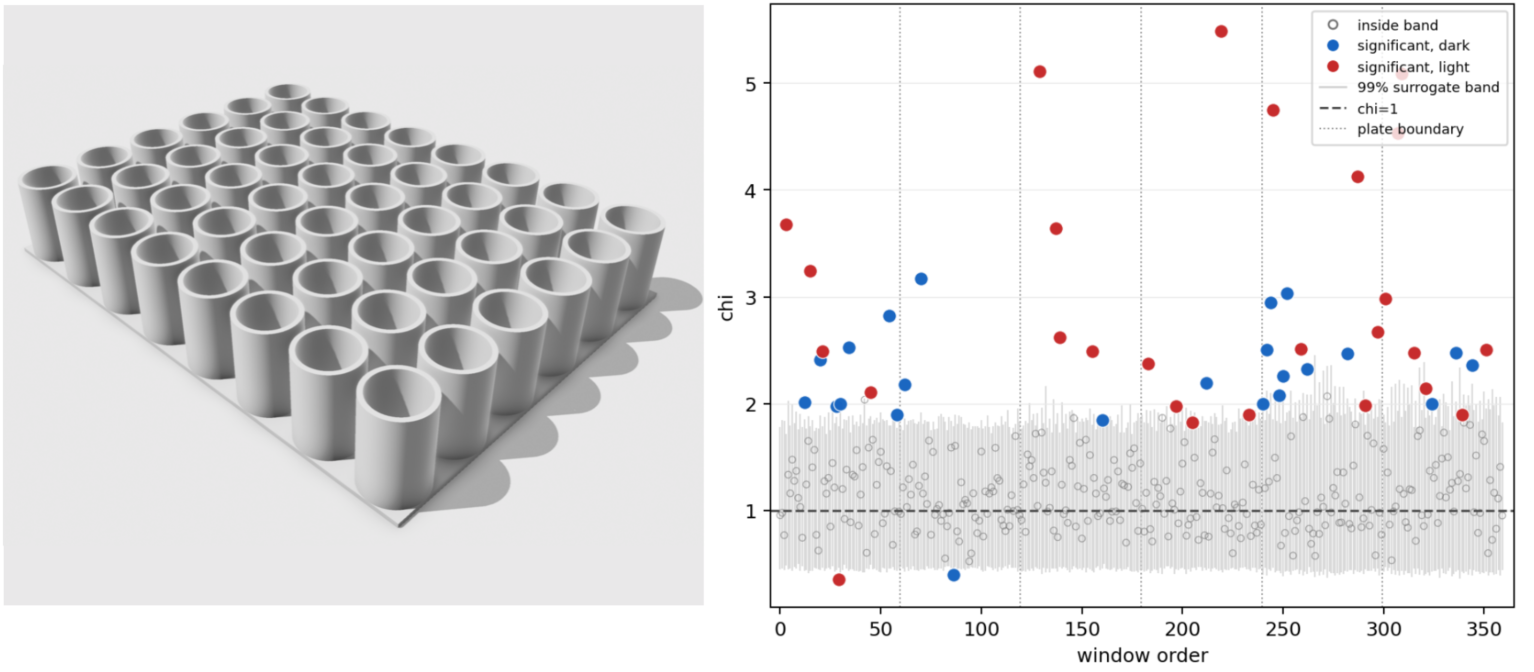
Left: 3D printed 48-well plate with isolated thin-walled wells separated by air gaps. Right: Common-mode coordination in isolated wells. Each point is the DIST-based chi statistic (χ = N·Var(M)) from the central 30 s window of one light or dark period, ordered by plate and then by time. Thin grey bars show the 99% circular-shift surrogate band for each window, the dashed horizontal line marks the null expectation (χ = 1), and dotted vertical lines separate plates. Open grey points fall within the surrogate band; filled blue points are significant dark-period windows and filled red points are significant light-period windows. See Methods for statistic details.

The most plausible signalling route between wells is substrate vibration: a moving fly drumming or pushing against the floor of its well could launch mechanical energy into the surrounding plastic, which a neighboring fly might in principle detect. The effect should scale with the amplitude of the movement that generates it. A fly that moves more should couple more energy into the substrate and, if this mechanism were important, should drive stronger well-to-well coordination. Windows in which flies are more active would be expected to show greater common-mode coordination. We tested this directly by plotting χ against the RMS movement in each central 30 s background window (Figures 4C and 7). The relationship is essentially flat across all three plate materials, with slopes near zero and R² ≤ 0.05. This is especially clear in the plastic plates, where the highest-RMS windows at the right of the plot show no increase in χ and remain close to baseline. Significant windows are not concentrated among the most active samples. Coordination and movement amplitude are therefore largely decoupled. For the background common-mode signal, this is not the pattern expected from an amplitude-scaled substrate-borne acoustic channel. It argues against fly-generated vibration as the carrier of that slow background signal, while leaving the transition-locked Δ signal to be judged by the within/cross transition comparison and the material tests.

**Figure 6.**
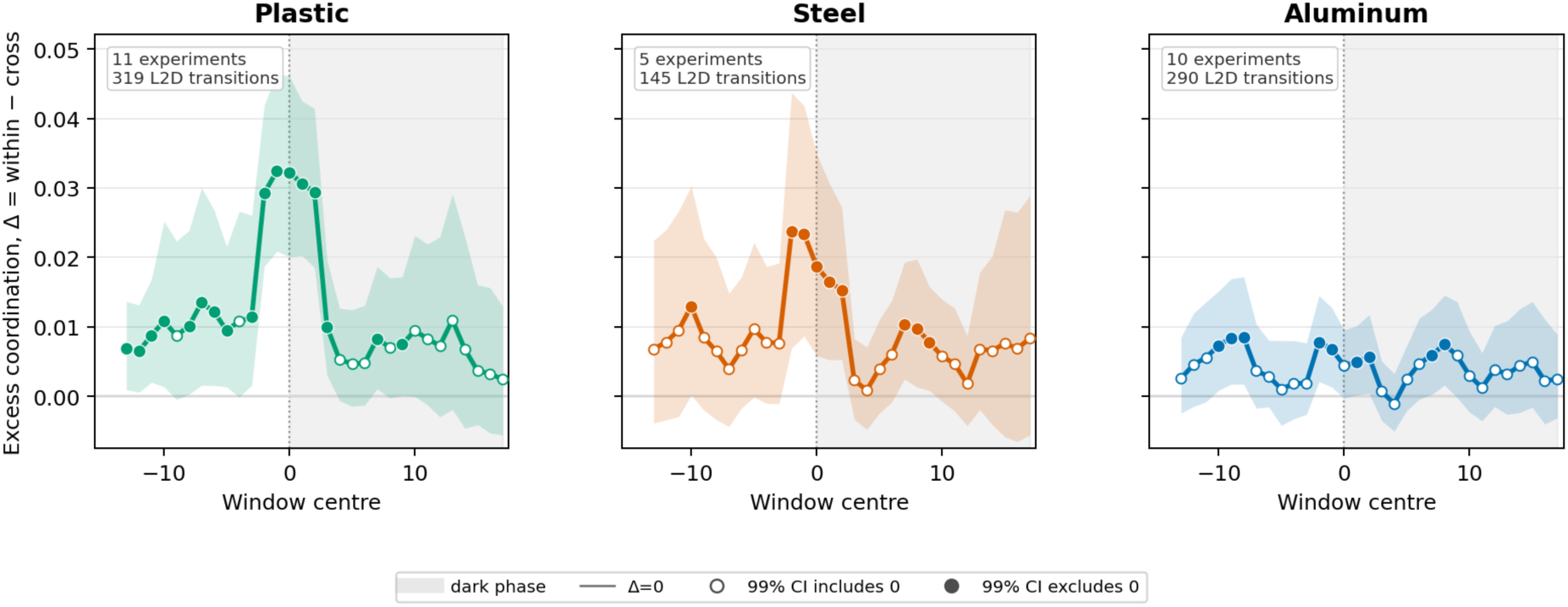
Continuous-displacement (DIST) excess pairwise coordination around the light-to-dark transition in three plate materials, ordered as plastic, steel, and aluminum. For each material, activity was analysed in 5-bin (5 s) sliding windows centred from –15 to +15 bins relative to the light-to-dark transition at 0. Lines show observed Δ = within-transition pairwise coordination minus cross-transition coordination; shaded bands are 99% moving-block bootstrap confidence intervals from 10,000 transition resamples. Filled points mark windows whose 99% CI excludes zero; open points mark windows whose CI includes zero. Grey shading indicates the dark phase after the transition. Window- and plate-level amplitudes for the material comparison are summarized in Supplementary Table S1.

**Figure 7.**
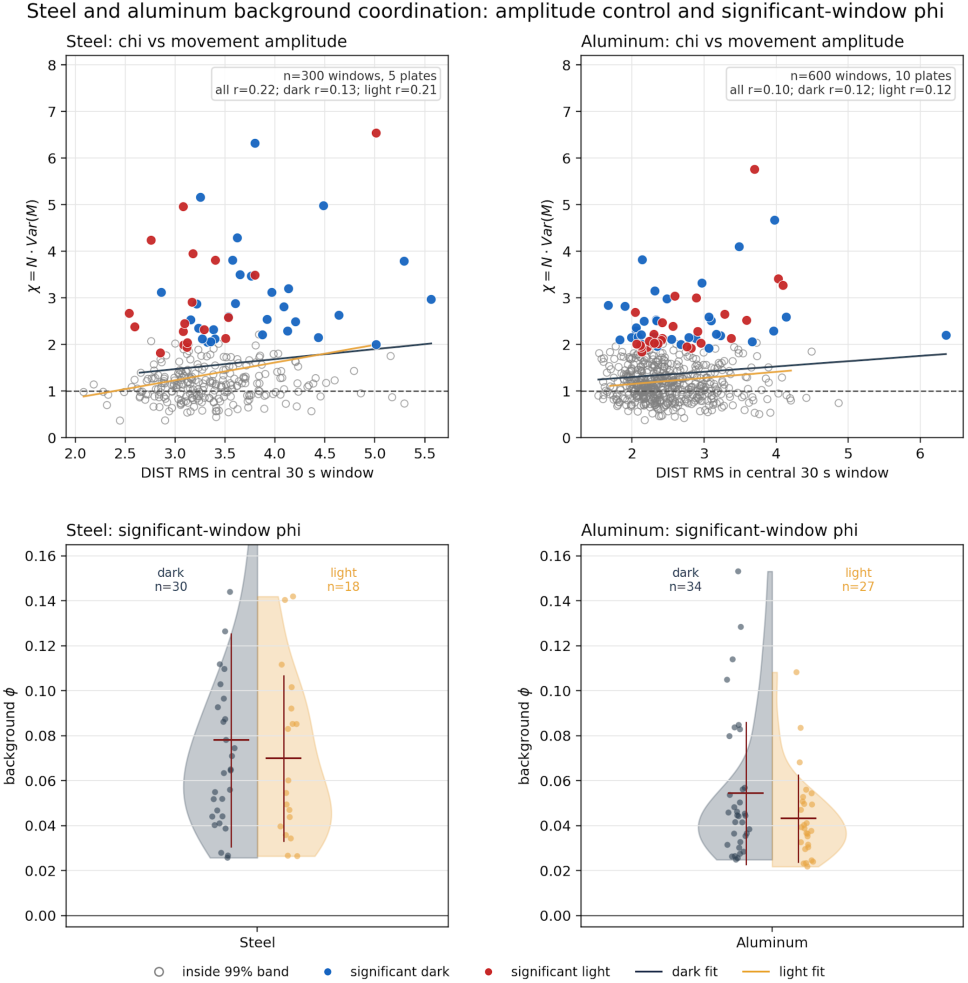
Steel and aluminum background common-mode coordination, movement amplitude, and significant-window φ. Upper panels: χ = N·Var(M) is plotted against DIST RMS for central 30 s light and dark background windows in steel and aluminum plates. Open grey points are windows within the 99% circular-shift surrogate band; filled blue and red points are significant high-χ windows in dark and light periods, respectively; colored lines show least-squares fits and dashed horizontal lines mark χ = 1. Lower panels: φ values for the significant high-chi windows in the same materials, split by dark and light periods. Dots are individual windows, half-violins show the distributions, and dark-red bars show mean ± SD.

**Figure 8.**
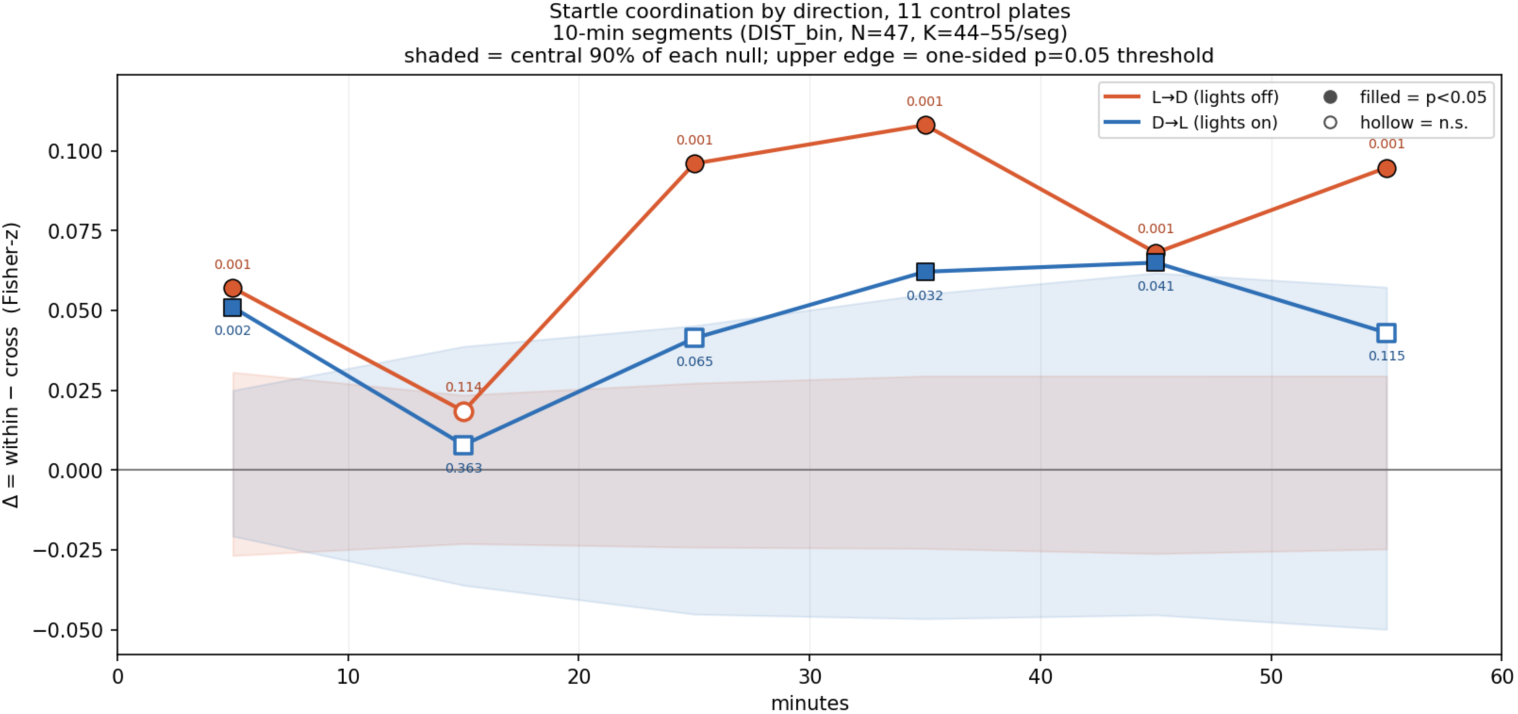
Transition-locked coordination in control plates is stronger for light*dark (lights-off) than dark*light (lights-on) transitions. Mean transition excess coordination Δ = ⟨z⟩within – ⟨z⟩cross across the N = 47 wells active in at least 9 of the 11 control plates, computed separately for light*dark ("lights off", orange) and dark*light ("lights on", blue) transitions and plotted for six consecutive 10-minute segments of the one-hour recording. For each transition, a 5-bin (5 s) window of binarised displacement beginning ≈2 bins before the transition was z-scored within each well; ⟨z⟩within is the mean pairwise Fisher-z Pearson correlation between wells taken within the same transition, averaged over the K = 44–55 transitions pooled across plates in that segment, and ⟨z⟩cross is the same correlation taken between wells in different transitions, which removes the component the wells share through common time-locking to the stimulus. Δ > 0 therefore reflects co-movement beyond the shared startle. Significance was assessed by an independent per-well permutation of transition labels (B = 1000 shuRes), which preserves each well’s activity distribution while destroying same-transition coupling; the value beside each marker is the one-sided permutation p = (1 + #{Δnull ≤ Δobs}) / (1 + B), with a floor of 0.001 at B = 1000. Filled markers denote p < 0.05, open markers non-significance. Shaded regions give the central 90% (5th to 95th percentile) of each null distribution, so their upper edge is the one-sided p = 0.05 threshold and a point clears its band precisely when p < 0.05. Both directions fall into the null at 10–20 minutes, the early non-stationary window seen across recordings; elsewhere L*D coordination is significant in 5 of 6 segments and D*L in 3 of 6, D*L running consistently weaker at roughly 0.6 times the L*D magnitude.

### Effect of well separation

To test inter-well coupling through the shared substrate directly, we 3D-printed a 48-well plate in which each thin-walled well was separated from its neighbors by an air gap, removing the continuous plastic that otherwise joins the wells. The aim was to break the structure-borne vibration path while leaving the electromagnetic path intact: an air gap is an acoustic impedance mismatch and will pass less structure-borne energy from one wall into the next, whereas both plastic and air are effectively transparent to a near-field magnetic signal. In six experiments the flies startled only weakly, but the background common-mode correlation was preserved, with χ exceeding the circular-shift surrogate band in many central light and dark windows much as in the solid plates. Quantitatively, mean χ was 1.31 in dark windows and 1.37 in light windows (180 windows each; circular-shift aggregate p = 0.0005 for both), with mean φ = 0.0088 and 0.0101 respectively and 25/180 dark and 26/180 light windows exceeding the one-sided 1% threshold. Coordination therefore persists when the continuous side-wall path between wells is interrupted by air gaps, although the wells still share a common imaging floor. This is diRcult for a purely structure-borne side-wall channel to account for and is the behavior expected if the carrier crosses the non-conducting gap as a near-field magnetic field. We also attempted the same recording in deaf *btv* (*beethoven*) mutants, in which auditory mechanotransduction is defective. These flies were too inactive to compute reliable coordination even in the background, so this experiment is an activity-failure null rather than evidence for or against the coordination mechanism.

### Coordination in metal plates

We have previously reported magnetic near-field RF emission from flies, using resonant cavities coupled to the magnetic rather than the electric component of the field^3^. This made near-field magnetic coupling a plausible candidate channel to test. At the millimeter distances between wells, any such coupling would be near-field: the short-range magnetic component of an oscillating electromagnetic field, rather than propagated waves. Conductive material between wells can weaken such fields because changing magnetic fields drive eddy currents in metal that oppose the original field. However, the comparison is not simply metal versus nonmetal: permeability, conductivity, geometry, and the transparent imaging aperture can all matter.

To test whether the coordination depended on plate material rather than simply on the presence of physical walls, we made plates with the same 48-well geometry from two metals and imaged them through the same transparent substrate. The two metals differ in the relevant electromagnetic properties: aluminum is a highly conductive, nonmagnetic metal, whereas mild steel is conductive and ferromagnetic. The coordination signal was absent in aluminum and approximately halved in mild steel. Signal amplitude therefore did not divide cleanly into plastic versus metal.

Figure 7 shows the relationship between background signal in the metal plates and RMS fly movement. The weak relationship between χ and movement amplitude in both metals, together with the plastic control in Figure 4C, argues against an amplitude-scaled substrate vibration explanation for the background common mode.

### Electrical conductivity vs magnetic permeability

The analysis in Figure 6 does not support a simple plastic-versus-metal interpretation: plastic is strongest, aluminum is weakest, and steel lies between. Steel and aluminum differ in at least two relevant ways. First, steel is 6-10 times less conductive than aluminum and eddy currents will be smaller. Second, steel is ferromagnetic, with a relative permeability of order 10²–10³, and can distort static and low-frequency magnetic fields by a mechanism distinct from eddy-current screening: its high permeability provides a low-reluctance path that draws magnetic flux into the walls and around the enclosed volume.

Regardless of material, metal wells cannot be expected to provide perfect RF shielding. The flies are not deep inside closed Faraday cavities; they sit close to the mouths of the holes, where fringing fields can leak around the conducting walls. Quantitatively this matters because near-field magnetic coupling scales approximately as 1/r³: moving the effective source or detector from 5–10 mm inside a well to within ∼1 mm of an aperture could increase leakage by roughly 10²–10³ relative to a recessed or closed geometry. The metal plates therefore test attenuation of inter-well coupling, not its absolute elimination. A decisive future experiment may require a transparent but conductive bottom plate, for example ITO-coated glass or a fine conductive mesh, so that flies can still be imaged while the aperture is electrically closed.

## Discussion

In summary, physically separated flies coordinate the magnitude of their movements around a shared light-dark startle more closely than the shared stimulus alone can explain. Their behavior also exhibits a weak background common mode coordination away from the L-D transitions. The two statistics give related but not identical material contrasts: the transition signal declines from plastic toward aluminum, whereas the background common mode is similar in plastic and steel and lower in aluminum, remaining significant in every material. Several of these observations could have a mundane explanation: the background common mode could reflect a shared plate-wide influence, the transition coordination could reflect some shared influence we have not identified, and the metal contrast could have an explanation we have missed. Two conventional signal sources seem unlikely: inter-well visual coupling is excluded by the opaque plate, and self-generated substrate vibration is disfavored because the most active windows do not show higher χ. Taken together with the fact that we have previously shown Drosophila emit spontaneous radiofrequency radiation that is powered by metabolism, tracks nervous system activity, and is abolished by anesthesia^3^, our results support a working hypothesis: flies emit and detect near-field magnetic RF fields. This remains a mechanism to test rather than one established by the present experiments.

If an emitted near-field magnetic RF signal is being detected, what detects it? The putative coupling is magnetic and oscillates at radiofrequencies, far too fast for any mechanical transducer to follow, and far too weak to exert a usable force on a bulk structure. Such a weak oscillating magnetic field carries negligible energy. One way for so weak an oscillating magnetic field to produce a chemical or behavioral change is through a system whose output depends on its effect on electron spin. Radical pairs are transient pairs of molecules, each with an unpaired electron whose spins are quantum-correlated, that interconvert between singlet and triplet states at rates that a weak resonant RF field can perturb, altering the yield of the downstream reaction^4,5^. This is the mechanism already invoked for magnetoreception and, to our knowledge, is the candidate most readily reconciled with the amplitude and frequency range of the signal we infer. We therefore regard a radical-pair sensor as the preferred hypothesis, and its identification by genetics as the central open problem.

If the detector is uncertain, the emitter is even more so. A known plausible source of radiofrequency emission from a biological system is the relaxation of electron or nuclear spins^6^: a spin flip is a change in magnetic moment and couples to the electromagnetic field magnetically. Known instances of electron-spin RF emission, the free-induction signal in pulsed ESR^7^, and chemically induced dynamic electron polarization (CIDEP)^8^ are also detected as near-field magnetic signals. We do not know what generates the radiofrequency signal, and our previous detection of it establishes its existence without identifying its source. One possibility is that a chirality-induced spin selectivity (CISS) effect is responsible. Electrons passing through chiral molecules (proteins, DNA) become spin-polarized in the direction of travel^9^, and the mitochondrial electron-transport chain drives large electron currents through exactly such chiral structures. Spins driven out of thermal equilibrium can in principle relax radiatively, emitting radiofrequency; this is the mechanism we previously proposed for the emission, and it links the signal to respiration and, through it, to neural activity.

### A speculative functional interpretation

The information content of such a signal might be low, perhaps closer to a global 1-bit pulse than to a complex message. If so, a conservative functional hypothesis is that the emission is for internal use, not communication, and that the coordination we observe is due to unavoidable near-field leakage. A nervous system that learns from reward must deliver to its synapses a signal that some recent action was worth repeating^10,11^. Dopamine supplies the established reward and punishment signals in the fly mushroom body, but it acts over chemical and circuit timescales. A brain-wide electromagnetic pulse, if it exists, would be fast and global by nature. On this speculative reading, dopamine would still set the sign and salience of reinforcement, while a metabolically generated electromagnetic emission could provide a rapid permissive signal to synapses poised to change.

If that is the emission’s purpose, the coordination we measured may simply reflect what happens when such an internal signal escapes the animal. Whether evolution makes any use of that is unknown. If it did, the leaked signal could provide a short-range cue about a neighbor’s current state or valuation of the environment, because a near-field emission falls off steeply with distance and would be useful only between immediate neighbors. This hypothesis is testable: the carrier frequency can be identified by perturbing coordination with synthesized near-field magnetic noise at defined bands; the sensor can be approached by genetic screening for loss of coordination in startle-competent flies; and the proposed internal role, reinforcement, can be tested by asking whether RF magnetic noise in the correct band disrupts learning. Here we report the coordination itself as evidence that nervous systems may be less electromagnetically private than is usually assumed.

## Methods

### Flies and husbandry

Experiments used Canton-S flies. The deafness control used *btv* mutants (y[1] w[*]; Mi{PT-GFSTF.0}CG5674 [MI08510-GFSTF.0] *btv*[MI08510-GFSTF.0-X]/CyO), which carry a defect in auditory mechanotransduction and were maintained in a yellow white (yw) background. Flies were reared on agar at 25 °C and 60–70% relative humidity under a 12:12 LD light cycle. Adults were tested at 2–5 days post-eclosion. A mixed population of males and females was used. Before loading, flies were anesthetized with CO2 for sorting and allowed to recover for at least 24 h before recording. The flies were loaded in 48-well plates containing 0.5 mL of food consisting of 2% agar, 5% sugar, and 0.25% nipagin. A single fly was loaded into each well. Recordings were performed during daylight hours.

### Behavioral apparatus and tracking

Locomotor activity was recorded using the Zantiks MWP Z2s system (Zantiks Ltd, UK), which images flies housed individually in the 48 wells of a multi-well plate arranged in an 8 × 6 grid. Illumination was internal from below. For the machined metal plates, the plate sat on a transparent substrate that formed the imaging floor, so flies were still illuminated and imaged through the substrate rather than through metal. This transparent optical path also means that the metal plates were not closed electromagnetic cavities; the imaging aperture is part of the same fringing-field geometry considered in the Results. Video was captured at 30 fps and reduced to per-well activity in 1 s bins. Three movement signals were derived from the tracking output and used throughout. DIST is the distance moved per 1 s bin. DIST_bin is its binarized form, scored as moved or not-moved using a threshold of zero. MSD is the mean-squared pixel change within the bin. These definitions are used wherever the three signals are referenced below.

### Plate fabrication and materials

Four plate types were used, all sharing the same 48-well layout. Wells were 11.9 mm in diameter, 20 mm deep, on a 13.5 mm center-to-center pitch. The opaque white plastic plate was 3D-printed on a Bambu P2S using PETG at 0.4 mm layer height, and opacity suRcient to exclude visual contact between wells. The aluminum and steel plates were machined to identical external geometry and well dimensions from 6061 aluminum and S235JR respectively, and were used sitting on a transparent substrate that provided the imaging floor. Thus metal surrounded the wells laterally, while the optical path through the floor remained open. The thin-wall plate was 3D-printed in PETG with each well formed as a tube of 0.5 mm wall thickness separated from its neighbors by an air gap of 1.1 mm; the tubes shared a common floor. This geometry was designed to interrupt the continuous solid side-wall path between wells while retaining the common imaging floor; both plastic and air are effectively transparent to a near-field magnetic signal, whereas an air gap is a poor conductor of structure-borne vibration.

### Light/dark protocol

Each recording ran for one hour under a protocol of interleaved light and dark periods of one minute each, giving 30 transitions of each type. The light-to-dark transition produces a sharp, reproducible startle across the plate and was the transition used for the transition-locked analysis. No lead-in period was used. The same protocol was applied across all plate materials. Isolated, unstimulated flies become quiescent, making it impossible to measure correlation in non-moving animals; the repeated startle creates high-activity windows where coordination can be resolved. The light-to-dark versus dark-to-light control is shown in Figure 8.

### Statistical analysis and reproducibility

Two levels of inference are distinguished throughout. Figure 1 summarizes the two coordination statistics used in the paper: Δ, a transition-locked excess after subtracting the shared startle, and χ, a background common-mode measure between startles. Window-level tests (the circular-shift transition null for Δ and the circular-shift background null for χ) are descriptive within a plate and localize candidate coordination. Top-level existence claims are made after collapsing each plate to one value, with the full plate-level analysis reported in Supplementary Materials. Because the analyses span three movement signals, two phases, three materials, and many windows, effect sizes and plate-level contrasts are emphasized over isolated window hits. Transition confidence intervals use 10,000 moving-block bootstrap resamples at the 99% percentile level. Analyses were performed in Python using scripts generated with Claude and GPT and then checked by inspecting the code, running synthetic and null controls where applicable, verifying transition counts and active-well masks against the raw workbooks, and reproducing the CSV summaries used for plotting. Data and analysis code will be deposited at Zenodo before submission. To quantify rather than assume the within-plate dependence that motivates the plate-level collapse, we measured the lag-1 autocorrelation of the per-transition Δ series and the per-window χ series within each plate, and converted it to an effective sample size under an AR(1) model, n_eff = n(1 – ρ4)/(1 + ρ4) (Figure 9). Serial dependence was weak (transition mean ρ4 = 0.13; background |ρ4| ≤ 0.18), so the pooled effective sample sizes, 584 of 754 transitions and 736–743 of 780 background windows, remain far above the plate count. This shows the window-level p-values are not materially inflated by serial correlation, but does not replace the plate-level tests, which additionally guard against a shared plate-wide offset that no autocorrelation correction can remove.

**Figure 9.**
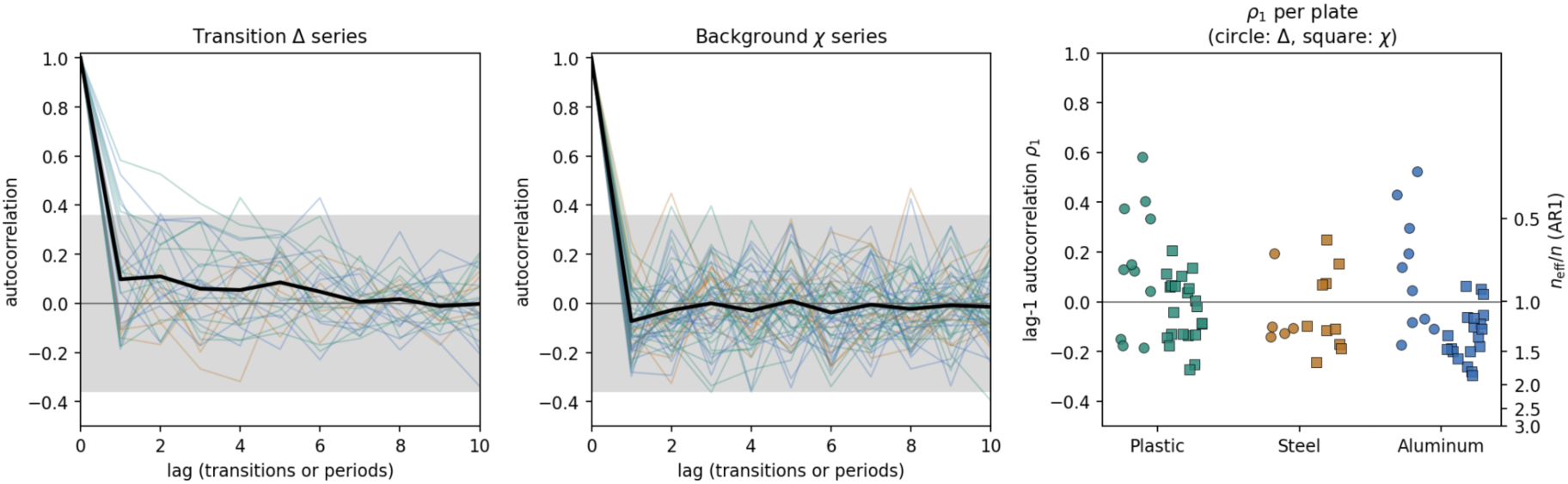
Within-plate serial dependence of the two coordination statistics. For every plate (11 plastic, 5 steel, 10 aluminum), the per-transition excess-coordination series (the within-transition Δ contribution, one value per light-to-dark transition in time order) and the background χ series (one value per central-30s light or dark period) were autocorrelated. Left and centre: autocorrelation functions per plate (faint, coloured by material) with the across-plate mean (black); grey band is the approximate 95% white-noise interval. The transition Δ series shows weak positive lag-1 autocorrelation (mean ρ4 = 0.13 across 26 plates) that decays to zero within a few transitions, as expected from slow arousal drift across the session; the χ series shows none (mean ρ4 between –0.04 and –0.18 depending on material and phase). Right: lag-1 autocorrelation per plate (circles, Δ; squares, χ) by material, with the implied AR(1) effective-sample-size fraction on the right axis. Summed over plates, the effective number of independent transitions is 584 of 754 (77%) and of background windows 743 of 780 in dark and 736 of 780 in light (94–95%). These greatly exceed the plate count of 26, indicating that the window-level analyses are not strongly pseudoreplicated.

### Transition-locked coordination (Δ)

For each light-to-dark transition, Δ was computed from all active well pairs in sliding windows of width W = 5 bins (5 s), with pairwise Pearson correlations Fisher-z-transformed before averaging. Window centres spanned –15 to +15 bins relative to the transition; wells with no variance in a window were excluded. Window-level significance used an autocorrelation-preserving circular-shift transition null (B = 10,000), in which each well’s ordered sequence of transition snippets was circularly shifted by an independently chosen offset. This preserves each well’s transition-to-transition structure and slow arousal drift while breaking simultaneous cross-well alignment. Confidence bands are 99% moving-block bootstrap intervals from 10,000 transition resamples. Window-level p-values localize candidate coordination in time; plate-level tests are reported in Supplementary Materials.

### Background common-mode statistic (χ and φ)

For background analyses, windows were the central 30 s of every light and dark period. Within each window, active wells were z-scored, M(t) was averaged across wells, χ = N·Var(M) was computed, and φ was recovered from χ = 1 + (N – 1)φ. Significance used a circular-shift null in which each well’s series was shifted by an independently chosen offset (B = 2,000), preserving each well’s autocorrelation and amplitude distribution while disrupting cross-well alignment. A window was counted as significant if χ exceeded the one-sided 1% threshold; reported summaries include mean φ across windows, φ within significant windows, and the fraction of windows passing that threshold. As a camera-artifact check, we processed one workbook recorded from dead flies. DIST contained no nonzero movement, so neither Δ nor χ was estimable; the run is therefore an activity-failure control showing no tracker-generated coordination signal, not evidence for or against biological coordination.

### Movement-amplitude control

A self-generated substrate channel should scale with the amplitude of the movement that drives it, so common-mode coordination should rise with per-window movement if vibration carries the effect. To test this, χ was plotted against the RMS of DIST computed in the same central 30 s window, separately for each material. A least-squares line was fitted and its slope and R² recorded; the test is whether the slope differs from zero and whether significant high-χ windows concentrate among the highest-RMS windows. Significant windows were identified as in the background analysis so that both analyses draw on the same windows.

### RF coupling measurement (VNA)

Inter-well radiofrequency coupling was measured as the two-port transmission S21 between two loop antennas connected to a vector network analyzer (Siglent SNA 5002A). The loops are electrically small and poorly matched and were not de-embedded; all values are therefore interpreted as matched-configuration differences rather than absolute coupling. S21 was swept from near DC to 4.5 GHz. Cable and connector reflections were removed by time-domain gating before each sweep. Measurements were made in two geometries: "close", with the loops lowered into neighboring wells in the geometry the flies occupy, and "far", with the loops maximally separated. Steel-minus-aluminum and close-minus-far differences were computed after applying identical processing to the paired spectra and are reported in dB. The close-versus-far comparison indicates a near-field contribution, while the steel-versus-aluminum comparison indicates material-dependent transmission under this probe geometry; because the probes can load and detune differently in the two metals, the measurement is not a de-embedded estimate of the field from a biological source.

### Data repository

Data and analysis code are available at 10.5281/zenodo.21486220

## Acknowledgments

LT thanks Ekin Daplan and Dan Girshovich for many useful discussions and Alex Charonitakis for tutorials on using Zantiks. Google Quantum AI and the Tiny Blue Dot Foundation are thanked for their support.

## Authors’ contributions

LT initiated project, LT, KD and ES designed experiments. KD performed experiments. LT analyzed data. LT wrote paper. KD, ES and LT revised paper.

## Supplementary Materials

Xenon anesthesia recovery

### Autocorrelation

Our transition and background analyses each aggregate hundreds of measurements (319 light-to-dark transitions in plastic, hundreds of central light and dark windows) and ask whether the resulting statistic exceeds what a null predicts. The null, and the confidence intervals built from it, treat every window as a fresh draw. But windows recorded seconds apart in the same plate are not fresh draws. A fly’s arousal drifts slowly through the hour, so its movement in one transition resembles its movement in the next. When samples are serially correlated in this way, the effective sample size, is smaller than the raw count.

### Plate-level inference

To avoid treating autocorrelated windows or repeated transitions as independent replicates, we repeated the key tests after collapsing each workbook to a single value and using plate as the unit of inference. The window-level circular-shift null and the 99% moving-block bootstrap confidence intervals remain in the figures as descriptive, within-plate timing analyses; they localize when coordination occurs but are not the top-level test for existence across plates.

For the background common-mode statistic, each plate was summarized by mean DIST χ and mean φ across the same central light/dark windows used in Figure 4. Plate-mean χ exceeded the independent-well expectation of 1 in every plate: plastic 11/11 plates, mean χ = 1.48 ± 0.20, one-sided sign p = 4.88 × 10–4; steel 5/5, mean χ = 1.45 ± 0.13, p = 0.031; aluminum 10/10, mean χ = 1.29 ± 0.12, p = 9.77 × 10–4. Plate-level exact permutation comparisons gave plastic > aluminum p = 0.0083, steel > aluminum p = 0.017, and plastic > steel p = 0.370 (one-sided). Plate-mean φ showed the same ordering (plastic 0.0162 ± 0.0064; steel 0.0172 ± 0.0056; aluminum 0.0103 ± 0.0052). Thus the background common-mode signal is plate-level reproducible in all three materials and is lower in aluminum than in plastic or steel, but not absent.

For the light-to-dark transition statistic, each plate was summarized by its mean DIST Δ over windows centered from -2 to +2 bins around the transition. Plastic showed plate-level excess coordination in all plates (11/11 positive; mean Δ = 0.0226 ± 0.0217; one-sided sign p = 4.88 x 10^-4^; Wilcoxon p = 4.88 x 10^-4^; one-sample t p = 0.0031). Steel was weaker and more variable than plastic but retained a positive transition-associated signal in 4/5 plates (mean Δ = 0.0187 ± 0.0265; sign p = 0.188); because only five steel plates were available, the sign test is underpowered for this material. Aluminum was the weakest material (7/10 positive; mean Δ = 0.00636 ± 0.0102; sign p = 0.172), consistent with no convincing transition-centered signal at plate level. An exact permutation comparison of transition-centered plate means gave plastic > aluminum (difference = 0.0162, one-sided p = 0.015), while plastic versus steel and steel versus aluminum were not resolved at plate level.

Window-level values are descriptive pooled-transition amplitudes from the 5-bin DIST sliding-window analysis; the CI is the 99% moving-block bootstrap interval. At centre –2, one-sided pooled-transition label-permutation tests (10,000 permutations) gave plastic > steel p = 0.228 and plastic > aluminum p = 1.0 × 10–4. These window-level p values are descriptive because transitions are nested within plates. Plate-level values are workbook means over window centres –2 to +2 and are the inferential unit. In the plate-level material comparison, only plastic exceeded aluminum (difference = 0.0162, one-sided exact permutation p = 0.015); plastic versus steel (p = 0.355) and steel versus aluminum were not resolved.

### Steel and aluminum differ in inter-well RF coupling

In order to get a qualitative understanding of the magnetic RF properties of the two materials, we compared radiofrequency transmission between adjacent wells directly. Two small loop antennas were connected to the two ports of a vector network analyzer, and the transmission S21 between them was recorded from near DC to 4.5 GHz (Figure S2); cable and connector reflections were removed by time-domain gating before each sweep. These measurements are interpreted as material-dependent near-field transmission indicators rather than absolute inter-well coupling values for a biological source. Measurements were made in two geometries: with the loops lowered into neighboring wells ("close") and with them maximally separated ("far"). We report steel-minus-aluminum S21 differences in dB for matched geometries, together with the close-versus-far comparison that supports a near-field contribution. In the close geometry, steel showed higher S21 than aluminum across much of the sub-gigahertz range, typically by several dB, with frequency-dependent reversals at higher frequencies. In other words, the steel wells appeared less isolated from one another than the aluminum wells at low frequencies under this probe geometry. This is not what bulk conductivity alone would predict, and is consistent with steel’s ferromagnetic permeability: at these frequencies a high relative permeability can provide a flux path that partly offsets eddy-current screening, whereas non-magnetic aluminum screens by eddy currents alone.

These supplementary figures collect the additional analyses generated during data processing that are not presented as main-text figures. The transition figures use the autocorrelation-preserving circular-shift null, and the confidence-interval figures use 99% moving-block bootstrap intervals as described in Methods. Exploratory controls are labelled as such.

---

**Figure S1.**
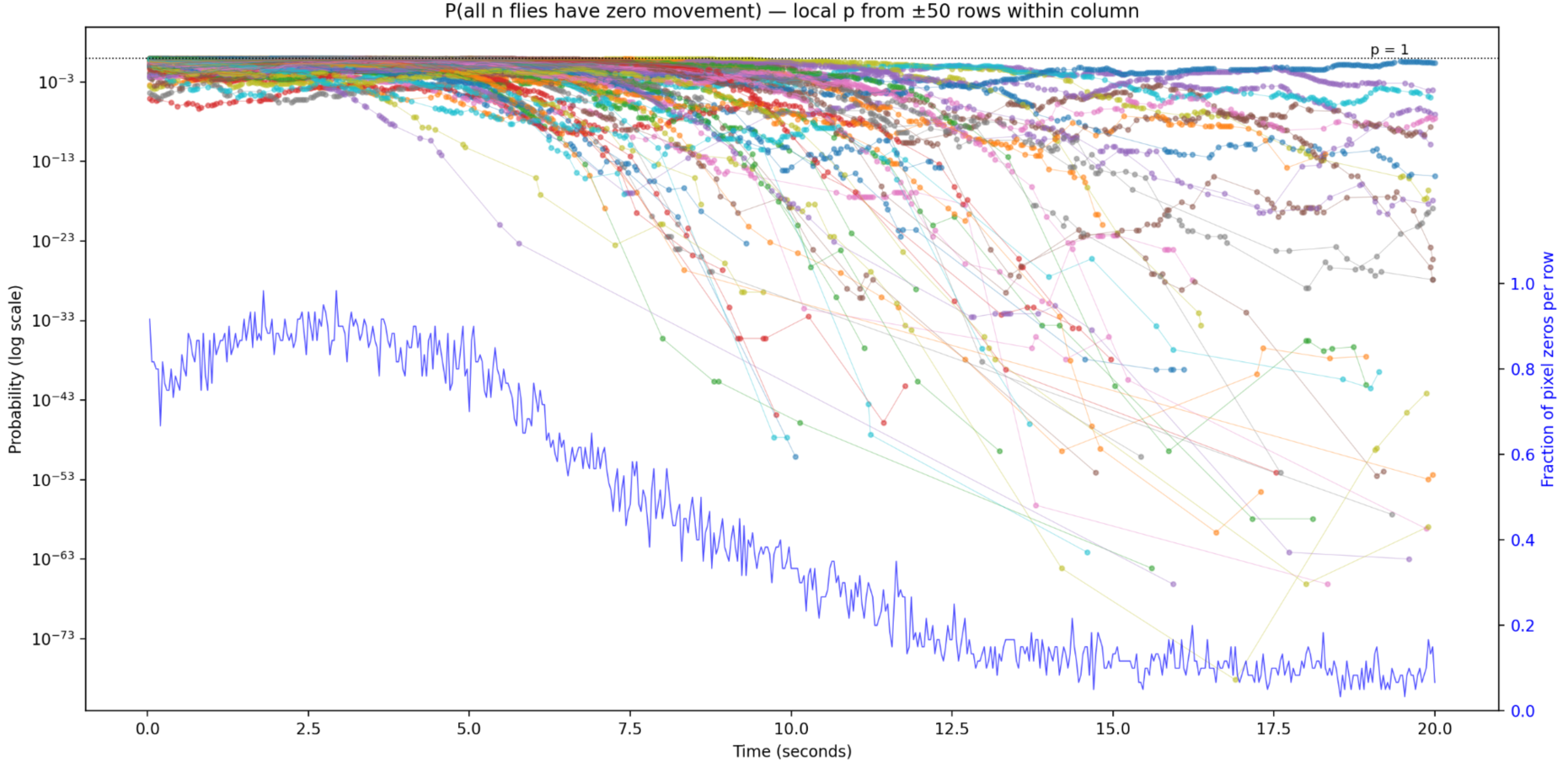
The probability that all tracked flies in a given video show zero total pixel movement simultaneously, plotted as a function of time (60 fps, every other frame). The method is described in Daplan and Turin, 2026. Forty flies were loaded per experiment, of which approximately 35 were successfully tracked in each video. At time zero all the flies are anesthetized, and the fraction of video frames showing them to be motionless is close to 1 (blue trace, right scale). At time zero the pressure in the chamber is gradually decreased from ≈2 atm to ambient over a period of 20 seconds and the flies wake up. The blue trace records the fraction p of frames in which the flies are found to be motionless (number of motionless frames per fifty frames). For each zero-valued cell in the pixel movement data (61 experiments in total), the number of flies n was obtained from the corresponding fly tracking data. The plotted quantity is p^n, representing the probability under the null hypothesis that flies move independently with identical stop probability p. Each color represents one of 61 videos. Unpublished data kindly shared by Dr Ekin Daplan.

**Figure S2.**
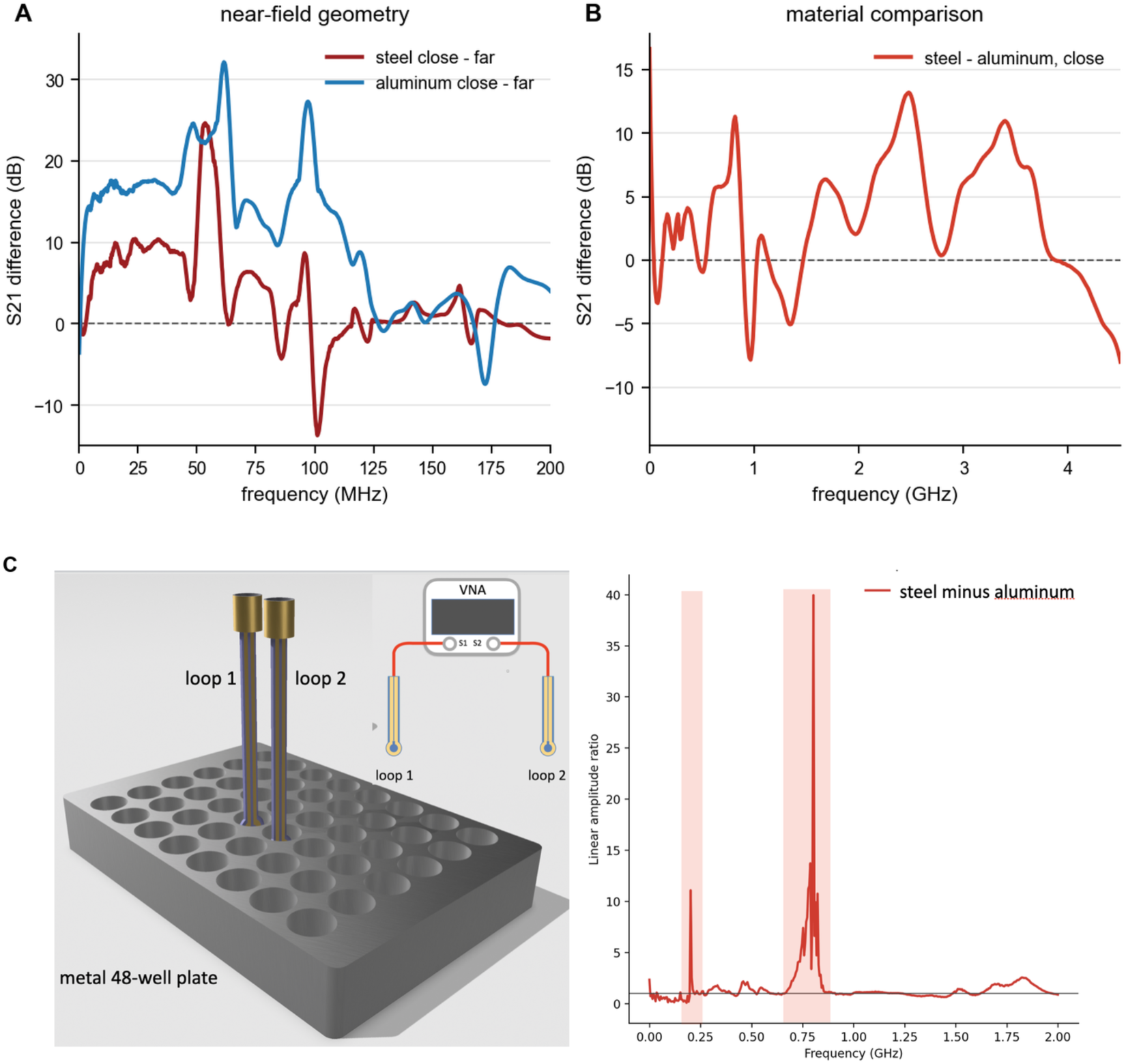
Material-dependent near-field transmission measured with loop probes. (A) Close-minus-far S21 difference for steel and aluminum plates, plotted in dB; positive values mean greater measured transmission in the close geometry than in the far geometry. (B) Steel-minus-aluminum S21 difference for the close geometry, plotted in dB after identical time-domain gating and smoothing; positive values mean higher measured transmission in steel than in aluminum. (C) The original display is restored as a separate panel: schematic of the VNA loop-probe geometry and the original linear-scale steel/aluminum close comparison, with shaded frequency regions marking the prominent peaks. Quantitative statements in the text refer to the dB S21 differences in A and B; C is retained as the original qualitative display. Because the loop probes are electrically small, poorly matched, and not de-embedded, these measurements indicate material-dependent near-field coupling but do not uniquely separate inter-well propagation from probe loading or detuning.

**Figure S3.**
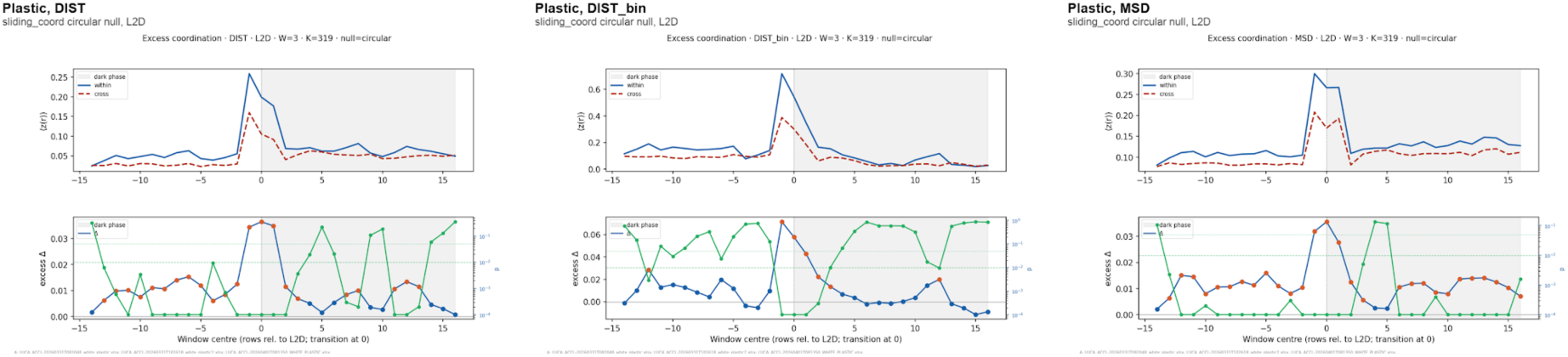
Formal circular-null transition analysis for white plastic plates. Each panel shows the light-to-dark sliding_coord output for one movement signal. The upper axis shows Fisher-z transformed within-transition and cross-transition pairwise correlations. The lower axis shows excess coordination, Δ = within - cross, together with window-level circular-shift p values from 10,000 permutations. These plots provide the formal p-value companion to the confidence-interval transition figure in the main text.

**Figure S4.**
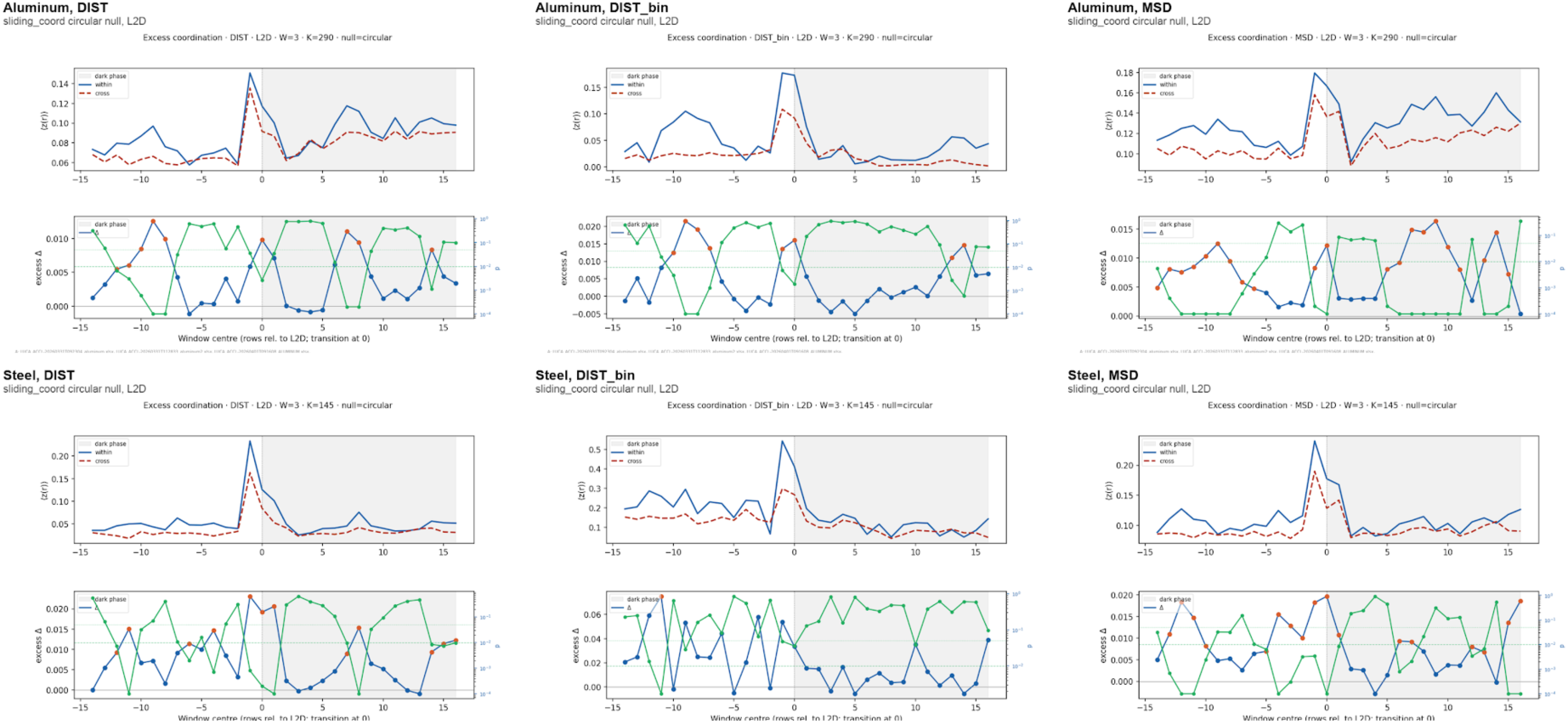
Formal circular-null transition analysis for steel and aluminum plates. Panels are arranged by material and movement signal, using the same statistic and permutation null as Figure S3. The metal-plate results show positive transition-associated windows in both materials, but with more fragmented support than in plastic, especially when judged as window-level evidence rather than as independent plate-level replication.

**Figure S5.**
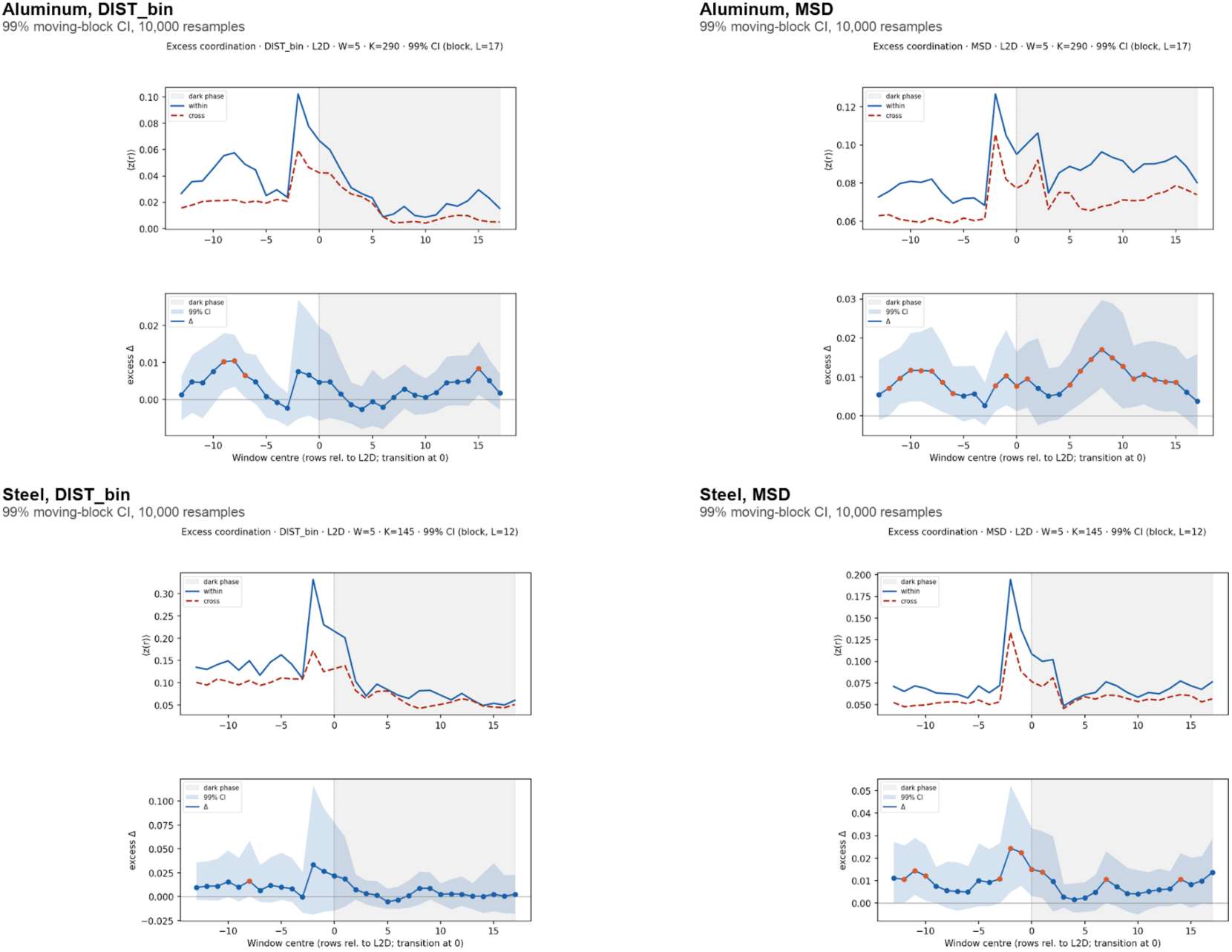
Metal-plate transition confidence intervals for DIST_bin and MSD. Steel and aluminum are shown for the two non-DIST movement signals using 99% moving-block bootstrap confidence intervals with 10,000 transition resamples. The figure complements the main-text DIST comparison by showing that the binary movement and mean-squared pixel change signals give related but noisier transition profiles in the metal plates.

**Figure S6.**
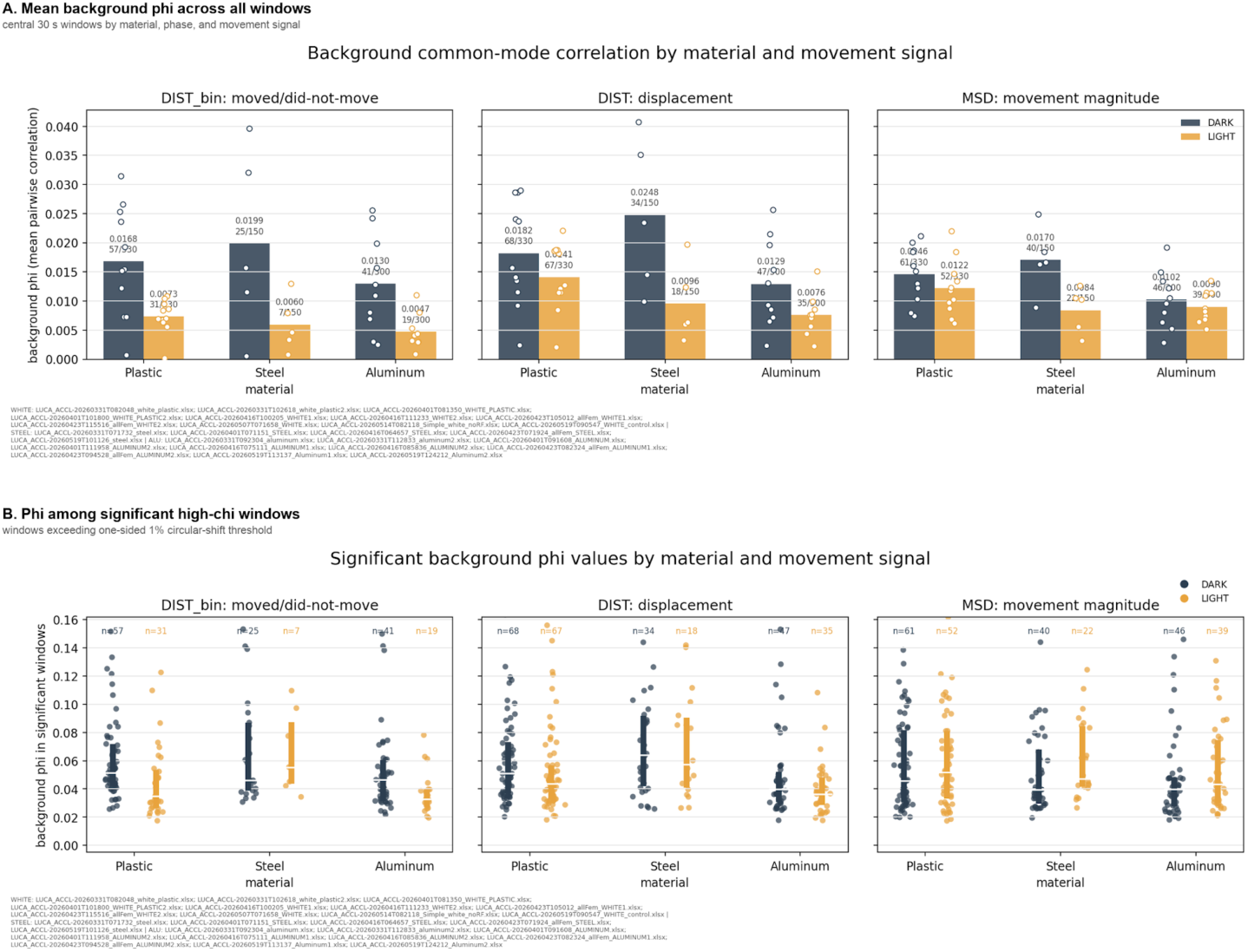
Background common-mode pairwise correlation across all movement signals. Panel A shows mean background φ in central 30 s light and dark windows for DIST_bin, DIST, and MSD across plastic, steel, and aluminum. Panel B shows φ restricted to windows whose χ exceeded the one-sided 1% circular-shift surrogate threshold. The figure extends the main-text DIST background analysis to the other movement signals.

**Figure S7.**
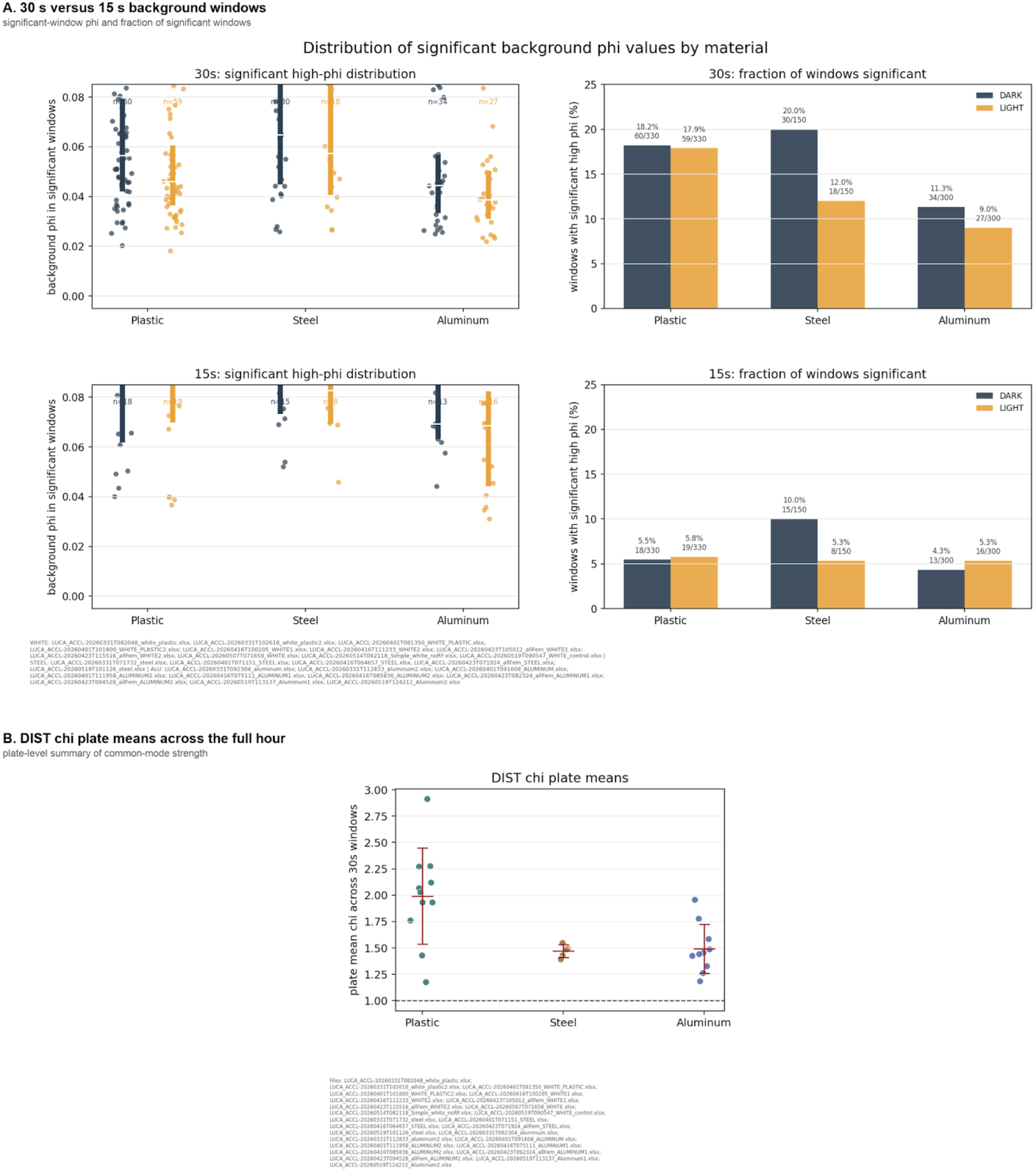
Robustness checks for the background common-mode statistic. Panel A compares 30 s and 15 s central-window definitions, showing that the material ordering and significant-window fractions are not artifacts of using the full 30 s middle window. Panel B is a broader robustness analysis that averages consecutive 30 s windows across the full hour, including transition-containing windows. These full-hour plate means are consequently higher than the transition-free central-background plate means used for inference and reported above.

**Table S1.** Light-to-dark transition amplitudes and comparisons with plastic at window and plate level.

| Material | Window $\Delta$ (99% CI) | Window p vs plastic | Plate $\Delta$ (mean ± SD); sign p | Plate p vs plastic |
| --- | --- | --- | --- | --- |
| Plastic | 0.0324 (0.0132 to 0.0562)<br>centre -1 | reference | 0.0226 ± 0.0217<br>(n = 11)<br>sign p = $4.88 \times 10^{-4}$ | reference |
| Steel | 0.0237 (-0.00329 to 0.0706)<br>centre -2 | 0.228 | 0.0187 ± 0.0265<br>(n = 5)<br>sign p = 0.188 | 0.355 |
| Aluminum | 0.00781 (0.000453 to 0.0183)<br>centre -2 | $1.0 \times 10^{-4}$ | 0.00636 ± 0.0102<br>(n = 10)<br>sign p = 0.172 | 0.015 |

## Notes

https://doi.org/10.5281/zenodo.21486220

### Competing Interest Statement

The authors have declared no competing interest.

